# Perturbed functional networks in Alzheimer’s Disease reveal opposing roles for TGIF and EGR3

**DOI:** 10.1101/286674

**Authors:** Saranya Canchi, Balaji Raao, Deborah Masliah, Sara Brin Rosenthal, Roman Sasik, Kathleen M. Fisch, Philip De Jager, David A. Bennett, Robert A. Rissman

## Abstract

While Alzheimer’s disease (AD) is the most prevalent cause of dementia, complex combinations of the underlying pathologies have led to evolved concepts in clinical and neuropathological criteria in the past decade. Pathological AD can be decomposed into subsets of individuals with significantly different antemortem cognitive decline rates. Using transcriptome as a proxy for functional state, we preselected 414 expression profiles of clinically and neuropathologically confirmed AD subjects and age matched non-demented controls sampled from a large community based neuropathological study. By combining brain tissue specific protein interactome with gene network, we identify functionally distinct composite clusters of genes which reveal extensive changes in expression levels in AD. The average global expression for clusters corresponding to synaptic transmission, metabolism, cell cycle, survival and immune response were downregulated while the upregulated cluster had a large set of uncharacterized pathways and processes that may constitute an AD specific phenotypic signature. We identified four master regulators across all clusters of differentially expressed genes by enrichment analysis including *TGIF1* and *EGR3.* These transcription factors have previously not been associated with AD and were validated in brain tissue samples from an independent AD cohort. We identify *TGIF1,* a transcriptional repressor as being neuroprotective in AD by activating co-repressors regulating genes critical for DNA repair, maintaining homeostasis and arresting cell cycle. In addition, we show that loss of *EGR3* regulation, mediates synaptic deficits by targeting the synaptic vesicle cycle. Collectively, our results highlight the utility of integrating protein interactions with gene perturbations to generate a comprehensive framework for characterizing the alterations in molecular network as applied to AD.

## 1. Introduction

Increase in aging population with improved longevity reinforces the urgency for prevention and treatment of progressive neurodegenerative diseases including Alzheimer’s disease (AD), the most common cause of dementia ^11, 41^. While, the predominant pathology of AD is accumulation of neuritic β-amyloid (Aβ) plaques and neurofibrillary tangles containing phosphorylated tau protein, co-occurrence of other neuropathological features is increasingly recognized to be a frequent event in brains of demented patients ^1, 11^. These changes including but not limited to inflammation, neuronal and synaptic loss, problems with blood circulation, and atrophy correlate with clinical symptoms of cognitive decline and have led to changes in diagnostic criteria during the last decade ^6, 41, 78^.

Multiple causal factors underlie the complexity of sporadic AD. Primary risk factors include age, gender and family history ^41^. The presence of elevated blood cholesterol, diabetes, depression, multiple lifestyle and dietary factors are also associated with increased risk of AD dementia, although not necessarily with AD pathology ^6^. Genetic mutations in amyloid precursor protein *(APP),* presenilin 1 *(PSEN1)*, and presenilin 2 *(PSEN2)* associated with autosomal dominant AD were critical in identifying pathogenic mechanisms associated with Aβ accumulation ^5, 83^. While many of the GWAS identified genes including the ε4 allele of apolipoprotein E (*APOE*) have implications of Aβ / tau processing, it is interesting to note that majority of the risk loci associate strongly with pathways involved in inflammation, lipid metabolism and endocytosis, which may partially explain the association of non-genetic risk factors ^46, 99^. Translation of molecular insights into predictive screening and diagnosis is important since clinical diagnosis of AD is difficult and often imprecise ^41^. This is complicated by the fact that genetic mutations explain only a small proportion of autosomal dominant AD and presence of *APOE* ε4 is not sufficient to cause AD ^6, 11, 41^. In addition, considering the variability in the rate of cognitive decline between individuals, transcriptomic changes at preclinical, prodromal and dementia stages of AD may be distinct and warrant subject selection refinement ^11, 95^.

In this study, we selected 414 clinically and neuropathologically confirmed AD subjects and cognitively normal age-matched controls, all sampled from a large well characterized community based neuropathological study ^6, 68^. Integration of gene perturbations with protein interactions is a powerful method to identify genes causal of a phenotype that are functionally cohesive, physically interact and share coherent biological pathways ^66^. We characterize the molecular network dysregulation in AD relative to controls by integrating global gene expression profiles with a pre-calculated brain tissue specific protein-protein interactome ^34^. Community detection applications to the human brain transcriptome have revealed intrinsic topological organization of functional co-expressed modules, they generally consider single interaction data ^46, 99 66^. By integrating the transcriptomic changes with protein interaction networks, we identify functional composite clusters by implementing the Louvain algorithm. The distinct gene clusters reveal extensive expression changes across multiple biological and cellular pathways implicated in AD along with novel candidates with less understood biological function. By applying the strategy of gene set enrichment, we identify four master transcriptional regulators across all the clusters. These include TGFB induced factor homeobox 1 (*TGIF1*), and early growth response 3 (*EGR3*), previously not associated with AD and validated by protein analysis of brain tissue samples from an independent AD cohort. We identify transcriptional repressor *TGIF* which modulates the disrupted *TGF-β* signaling, as being neuroprotective in AD by activating co-repressors regulating genes critical for arresting cell cycle, facilitating DNA-repair and restoring homeostasis ^64, 89^. We show loss of regulation of *EGR3* which is crucial for short term memory, to mediate synaptic deficits by targeting the synaptic vesicle cycle ^69, 78^.

## 2. Results and Discussion

### 2.1 Refined AD phenotype reveals perturbation in the transcriptomic landscape

After adjusting for covariates and multiple testing, total of 1722 genes were significantly differentially expressed in AD compared to age-matched non-demented controls (NDC) and 57% of those genes were downregulated (Figure 1A, Supplementary Table S1). Gene Set Enrichment Analysis (GSEA) ^82^ revealed functional enrichment of specific biological processes (GO) and molecular pathways (KEGG) underlying these genes. A total of 453 upregulated and 113 downregulated MSigDB gene sets ^82^ were identified at 5 % FDR. Among the GO terms enriched, those related to neural development, gliogenesis, metabolism and localization of proteins and extracellular structure organization were upregulated while those related to synaptic transmission, mitochondrial and metabolic processes as well as extracellular transport were downregulated (Supplementary Tables S2, S3). These results are consistent with previous findings of multiple studies associating AD pathophysiology to genetic perturbation and demonstrate the complexity of the disease ^11, 46, 68, 95, 99^.

**Figure 1.**
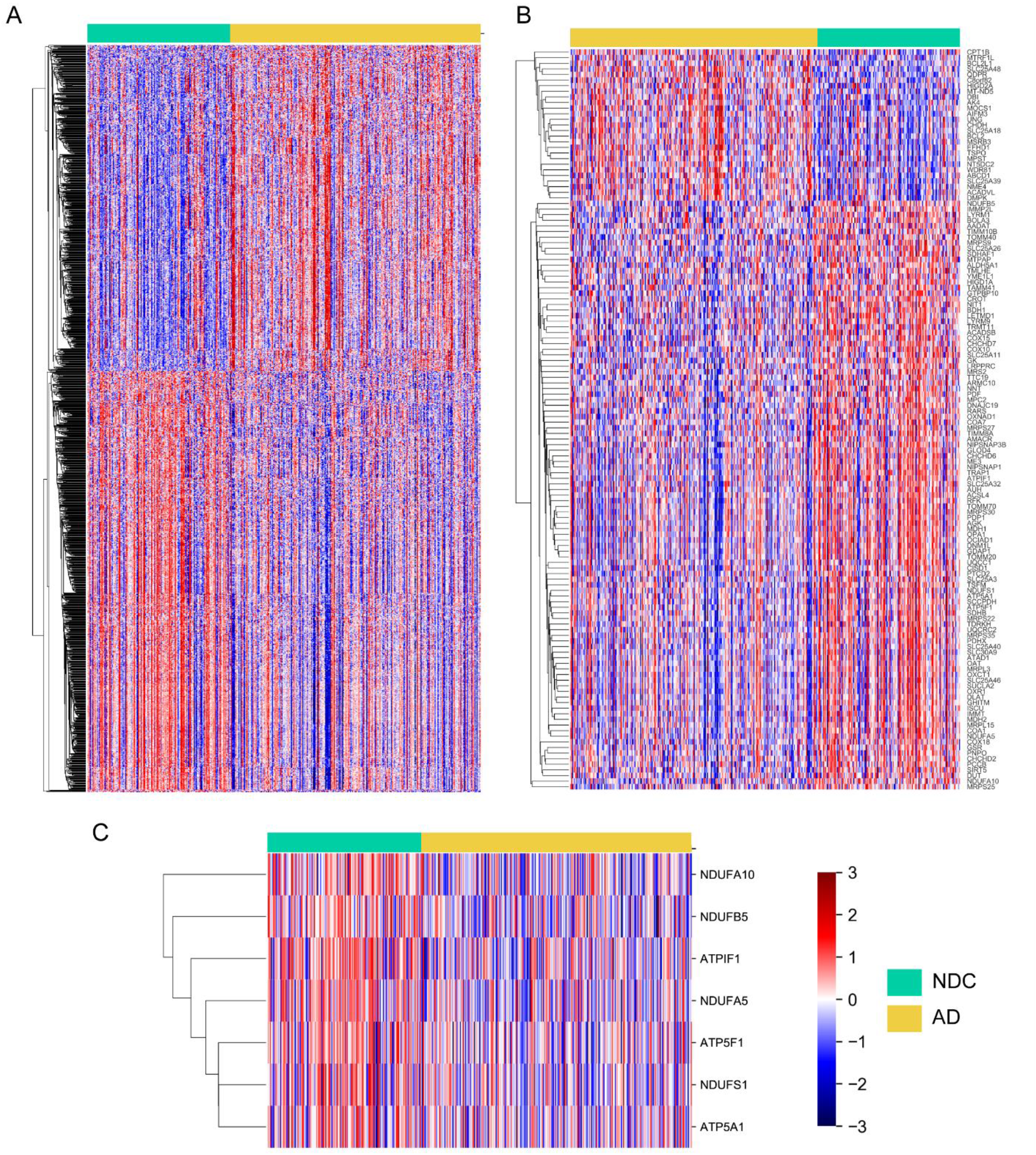

A total of 133 differentially expressed genes were also mitochondrial genes, defined as mitochondrially encoded genes from MitoCarta 2.0 ^16^ (Figure 1B) (Supplementary Table S4). Among these are nuclear-encoded genes for oxidative phosphorylation (OXPHOS) (Figure 1C). These include NADH ubiquinone oxidoreductase core subunits S1 (*NDUFS1*, PEP = 2.4e-02), A5 (*NDUFA5*, PEP = 8.0e-05), A10 (*NDUFA10*, PEP = 1.3e-02) and B5 (*NDUFB5*, PEP = 2.4e-02), all part of complex I involved in transfer of electrons to the respiratory chain. Reduced expression of complex V genes including ATPase inhibitory factor I (*ATPIF1*, PEP = 2.4e-03), subunit B (*ATP5F1*, PEP = 1.8e-03) and F1 alpha subunit (*ATP5A1*, PEP = 3.7e-02) have implications for production of ATP from ADP ^18^. In addition, expression of eight mitochondrial ribosomal proteins subunits (*MRP*) involved in translation of the mitochondrial-encoded OXPHOS genes are downregulated. Downregulated nuclear and mitochondrial genes encoding subunits involved in OXPHOS have been shown in the brain and blood of subjects with AD and mild cognitive impairment (MCI) compared to NDC ^25, 58^. While most mitochondrial genes were downregulated, 27 genes showed upregulation (Figure 1B). Of these, increased expression of diazepam-binding inhibitor (*DBI*, PEP = 8.9e-04), known for its role as mediator in corticotropin-dependent adrenal steroidogenesis and in modulation of the action of the GABA receptor has been shown to be dose dependent on Aβ aggregates^57^.

### 2.2 Aggregate network of protein and genetic interactions reveals novel gene candidates in biologically distinct clusters associated with AD pathogenesis

When integrated into the GIANT brain-specific interactome ^34^, the differentially expressed genes display distinct clustering structure (Figure 2A). These clusters were identified computationally using the Louvain modularity maximization algorithm ^88^. This graph-based unsupervised clustering analysis does not require any explicit assumptions, and instead uses an iterative process to optimally partition nodes in a graph, by maximizing the number of edges within clusters, and minimizing the number of edges between clusters.

**Figure 2.**
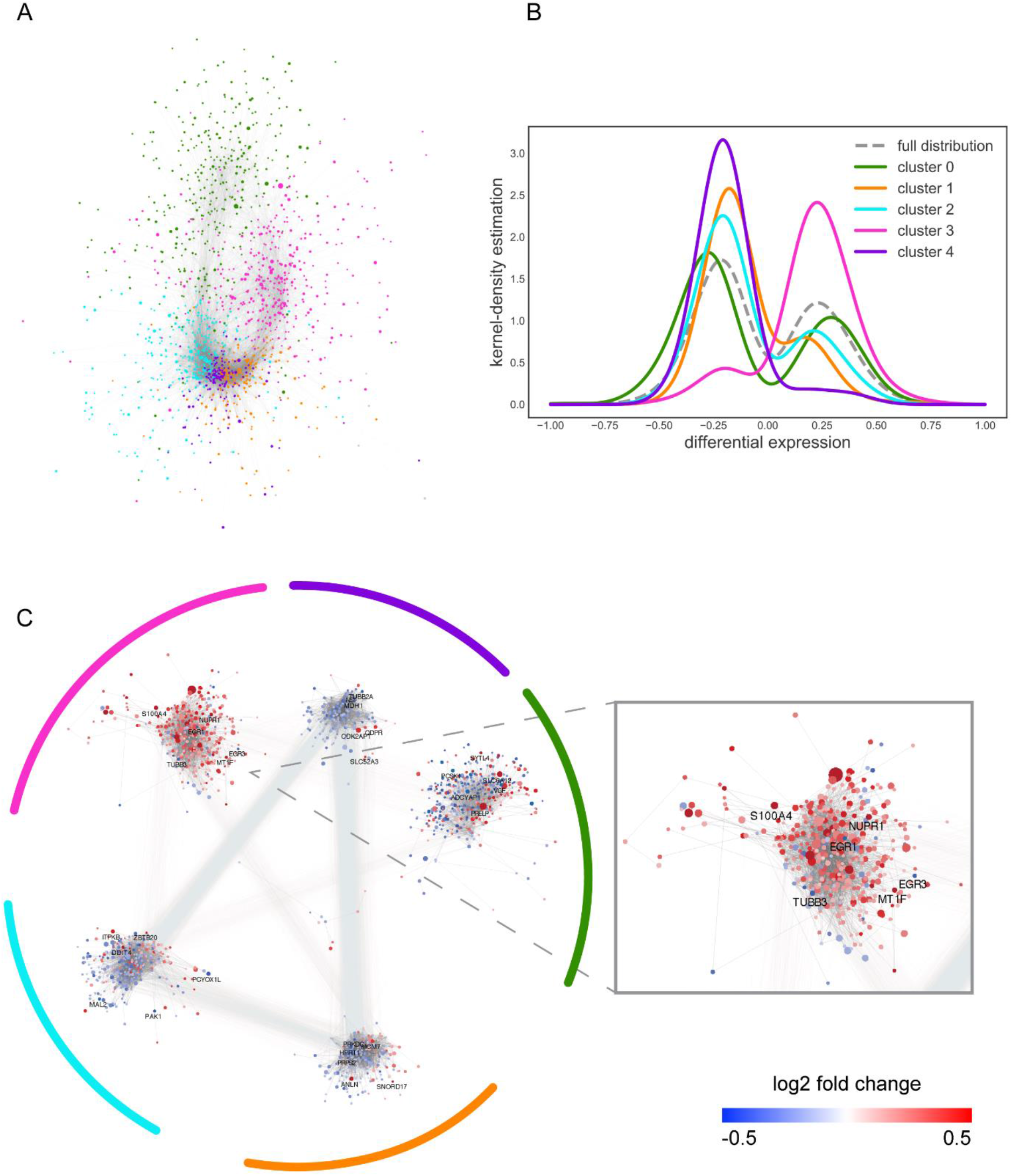

The largest connected subnetwork of 1391 differentially expressed genes partitions into 8 distinct clusters ranging in size from 2 to 373 genes. We restricted our analysis to clusters containing at least 5 genes and resulted in a subnetwork with 1382 genes. Many of these clusters are significantly up-regulated or down-regulated in AD with respect to NDC (Figure 2B, Supplementary Table S5). Two metrics were used to assess significance of up and down-regulation. The first metric was a simple binomial test, comparing the number of observed positive edges within each cluster to the number expected given the rate of positive edges within the full AD subnetwork. The second metric is the Kolmogorov-Smirnov test to check if the distribution of log fold change expression values in each cluster is significantly different than the distribution in the full AD subnetwork (Figure 2B). Of the 8 detected clusters, 5 that were enriched for up or down regulated genes are highly statistically significant (Figure 2C, Supplementary Tables S5, S6).

Overrepresentation analysis of the genes in the clusters characterized significantly enriched biological processes and molecular pathways. This revealed functionally distinct units related to broad categories of synaptic transmission, signal transduction, cell survival and viability, immune response and metabolism (Figure 3, Supplementary Table S7, Figure S1). Interestingly, the expression for clusters corresponding to synaptic transmission, DNA repair, immune response and metabolism were downregulated, while the cluster with overall upregulation had many uncharacterized genes (Figure 2C).

**Figure 3.**
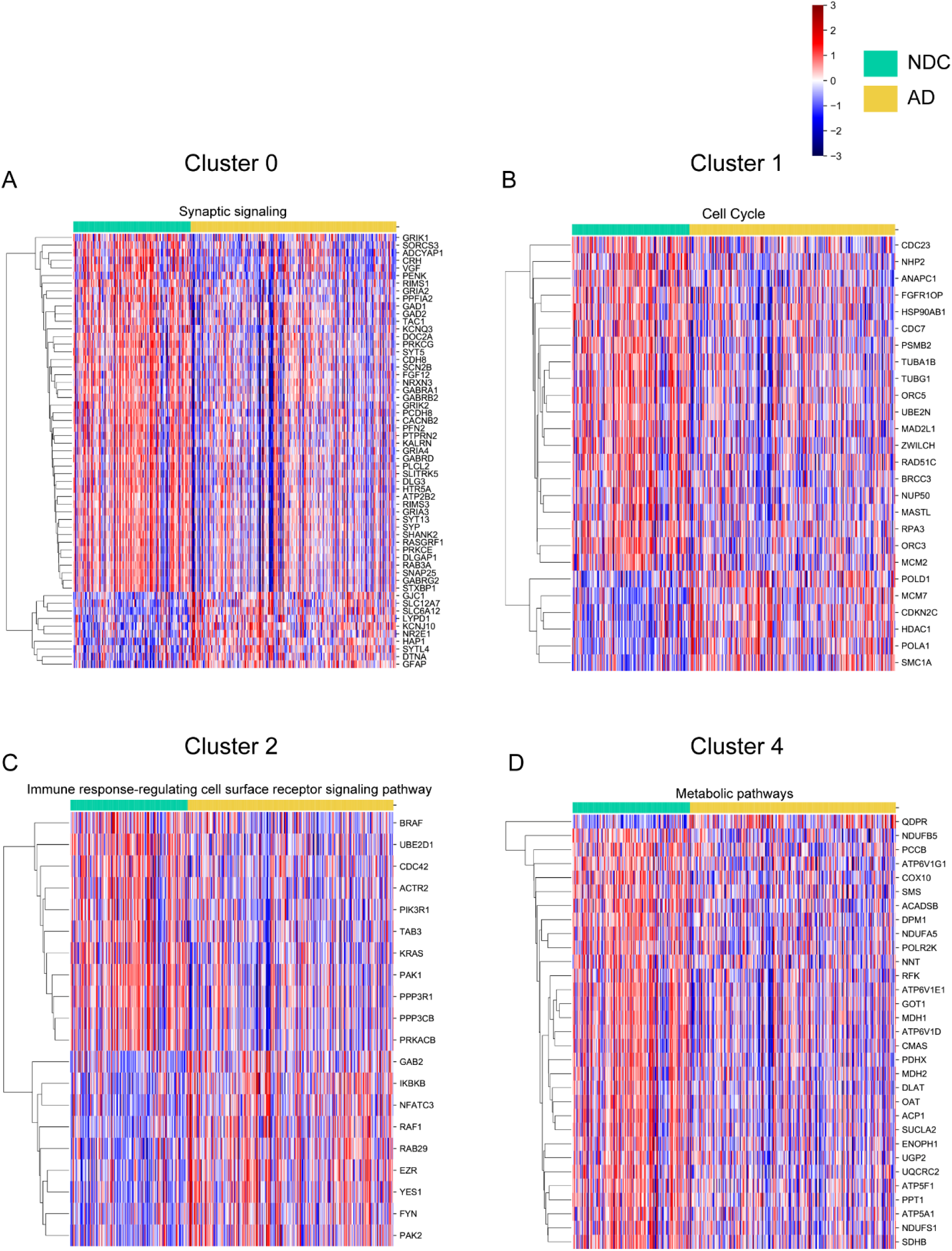

#### 2.2.1 Cluster0 – Synaptic Transmission

Loss of neurons and synapses in the hippocampus and cerebral cortex, a characteristic of AD is shown to strongly correlate with cognitive impairment, high level of Aβ production along with tau oligomers ^78^. Pathways in cluster0 were enriched for neurotransmitter related signaling (Figure 3, Supplemental Table S7, Figure S2). Expression of neuregulin-1 (*NRG1*, PEP = 6.7e-04) and neurexin 3 (*NRXN3*, PEP = 4.6e-03) which promote the formation of functional synaptic structures were downregulated ^63^. Expression for GABAA receptors was downregulated and is comparable to recent evidence of decreased amplitude of GABA currents on examination of the electrophysiological activity of AD brains which correlated with reduced mRNA and protein expression of α1 and γ2 GABA receptor subunits in the temporal cortex ^55, 57^. Downregulation in glutamate ionotropic receptor AMPA subunits (*GRIA2, GRIA3, GRIA4*) and kainate type subunits (*GRIK1, GRIK2*) along with *SHANK2* which forms a postsynaptic scaffold for receptor complexes is consistent with the idea that excess Aβ can enhance endocytic internalization of AMPA and NMDA receptors and suppress long term potentiation (LTP) in AD ^78^. Interestingly, increased expression of lysophosphatidic acid receptor 1 (*LPAR1,* PEP = 4.4e-03) a G-protein coupled receptor has been shown to regulate glutamatergic and GABAergic signaling although the details are unclear ^98^.

In addition to neurotransmitters, neurosecretory proteins and neuropeptides are well known for their role in neuronal cell communication. Reduced expression of neurosecretory protein VGF nerve growth factor inducible (*VGF*, PEP = 2.9e-05) considered to be associated with synaptic plasticity and function, has recently been identified in CSF and parietal cortex of AD patients ^21, 38^. These results confirm and extend the trend to prefrontal cortex in AD. Based on the specificity of VGF to the central nervous system, it is an ideal candidate with the potential for blood based biomarker for AD. Reduced cortical corticotropin-releasing factor immunoreactivity (*CRH*, PEP = 2.1e-04) in the face of increased hypothalamic expression) is a prominent neurochemical change in AD and is shown to mediate stress induced hyperphosphorylated tau through its receptor1 (*CRFR1*) with antagonist reducing Aβ levels in animal AD models ^71, 100^. Decreased expression of doublecortin like kinase 1 (*DCLK1,* PEP = 7.7e-03) which has both microtubule-polymerizing activity and protein kinase activity likely has implications for axon trafficking deficits ^49^. Interestingly, downregulation of dipeptidyl-peptidase 6 (*DPP6,* PEP = 1.5e-02) is associated with reduced dendritic spine density, a phenotype observed in AD brains ^56^.

#### 2.2.2 Cluster1 – DNA Repair and Transcription

Abnormal cell cycle reentry in response to accumulation of damaged DNA is hypothesized to precede AD pathogenesis ^23, 94^. Pathways in cluster1 were enriched for DNA replication and transcription (Figure 3, Supplemental Table S7, Figure S3) with approximately 15% of genes associated with cell regulated DNA repair and chromatin remodeling ^67^. One of the highly connected genes in this cluster, minichromosome maintenance complex component 7 (*MCM7*, PEP = 9.3e-05), was identified as an additional AD susceptible locus in large scale GWAS study ^27^. Expression of DNA-dependent protein kinase (*PRKDC*, PEP = 9.5e-03) which is essential for the repair of the double strand breaks, a lethal form of DNA damage was downregulated and is comparable to protein expression observed in AD brains ^17, 23^. Reduction in the base excision repair, a multistep DNA repair pathway has been detected in early stages of the disease (amnestic MCI) with continued trend with progression to AD ^94^. Reduced expression of tyrosyl-DNA phosphodiesterase 1 (*TDP1*, PEP = 4.0e-02), a DNA repair protein involved in base excision repair is a rational result ^26^. Overexpression of DEK Proto-Oncogene (*DEK*, PEP = 4.1e-02) known for its role in p53 destabilization, is associated with multiple cancer phenotypes and has not been characterized in AD with potential as an early target for neuronal homeostasis ^28^. PHD Finger Protein 19 (*PHF19*, PEP = 3.1e-05), a Polycomb-like protein is known to specifically bind to histone H3K36me3 and crucial for *PRC2* recruitment to CpG islands which mainly mediate transcriptional repression ^15^. Although its role in AD remain unclear, overexpression of *PHF19* as observed here has been shown to correlate positively with astrocytoma grades ^53^. Whether this was a consequence of cell state activity or an essential event is to be determined. Although not directly linked to AD, suppression of chromatin assembly factor I (*CHAF1B*, PEP = 2.0e-03) has recently been shown regulate somatic cell identity in a transcription factor-induced cell fate transition, correlated to protein levels in Down Syndrome brains and identified via whole exome sequencing to be associated with neurogenetic disorders with intellectual disability ^2, 19^. Interestingly, stathmin (*STMN1*, PEP = 3.3e-02) known to negatively correlate with neurofibrillary tangles and highly connected to genes in this cluster is shown to mediate cell cycle regulation via *MELK* kinase and has been investigated as a biomarker of DNA damage in AD ^61, 92^. This could provide an important link between dysregulation of microtubule polymerization and cell cycle dynamics as seen in AD and merits further investigation.

#### 2.2.3 Cluster2 – Immune Response

Immune system pathways were enriched in cluster2 (Figure 3, Supplemental Table S7, Figure S4). Several lines of evidence suggest that chronic neuroinflammation, shown to correlate with disease progression could contribute to the pathogenesis of AD ^46, 62, 99^. The associated causal regulators including *TYROBP* as described in Zhang et al ^99^ were not significant, although pathways under the ‘Fc’ and ‘Complement’ immune and microglia modules are enriched. While the role of C-type lectin receptors (CLR), members of the pattern recognition receptor family in AD are currently unclear, two functional variants of Mannose-binding lectin (*MBL2*), a soluble CLR are associated with AD risk ^80^. Antagonists to *MBL2* have shown to facilitate favorable outcome in experimental models of traumatic brain and spinal injury suggesting a potential target for modulation of immune response ^24, 33^. Increased expression of the chemokine receptor 4 (*CXCR4*, PEP = 3e-0.2) is shown to be associated with higher levels of activated phosphorylated protein kinase C and with synaptic pruning in early development ^65, 93^. Interestingly, this was concomitant with decreased expression of C2H2 zinc finger protein (*PLAGL1*, PEP = 4.2e-07) known to activate SOCS3, a suppressor of cytokine signaling ^76^. Fc receptors mediated glial cell activation identified as one of the immune pathways in AD GWAS study, has been attributed in the adverse side effects associated with the failed AD immunotherapy trials reiterating the need to understand the functional roles of these receptors ^32, 46^. Decreased expression of bromodomain family member (*BRWD1*, PEP = 1.7e-06), a histone reader essential for B lymphopoiesis is in lieu with the observed loss of the cross talk between innate immune system of the brain and the adaptive immune system as facilitated by circulating B and T cells ^59, 62^.

#### 2.2.4 Cluster3 – Novel Gene Candidates

Genes from cluster3, the only cluster with an overall upregulation in the differential expression were not readily characterized. No enriched pathways were detected, and only a few marginally significant GO terms were found. Examination of this cluster revealed more uncharacterized genes than expected by chance (p < 0.05), suggesting genes within this cluster would be attractive candidates for follow up studies as many encode for proteins not previously characterized or associated with AD. Enriched biological processes included small GTPase mediated signal transduction (FDR = 6.21 e-05), which are known to control diverse cellular activities (Supplemental Figures, S1). Evidence suggests that dysregulation of Rho-GTPase specifically *RHOA*, *RAC1* and *CDC42* mediated actin dynamics could be a key contributor to synaptic deficits observed in AD, yet interactions between the different signaling pathways remain unclear ^40^. Ras homolog family members C (*RHOC*, PEP = 4.5e-02), D (*RHOD*, PEP = 9.5e-03), G (*RHOG*, PEP = 1.2e-03) were upregulated and increased expression of ras-associated protein rab13 (*RAB13*, PEP = 4.9e-03) paralog for RAB8B is consistent with the protein levels measured in AD brains ^40^. ADP-ribosylation factor (*ARF3, ARF5*) sub family of GTPase have critical roles in secretory pathway for intracellular ER-golgi trafficking and in vesicle formation ^40^. *ARF6*, a paralog of *ARF3* was reported to be an important modulator of *BACE1* sorting in early endosomes in a clathrin-independent route and abrogating its function led to increased Aβ secretion while functions of *ARF5* in AD are unknown ^74^. AD GWAS identified gene families including *INPP5D*, *NME4, ABCA2, PLXNB1* were upregulated ^99^. Interestingly, transcription factor EB (*TFEB*) a master regulator for lysosomal biogenesis was overexpressed which is associated with regulated autophagy ^77^. Given that the events in AD are non-linear, whether the observed changes are pathogenic or protective is to be determined.

#### 2.2.5 Cluster4 – Metabolism and Bioenergetics

Metabolic dysfunction is a well-established characteristic of AD and pathways in cluster4 were enriched for energy metabolism and vesicle mediated transport (Figure 3, Supplemental Table S7, Figure S5). Clinical imaging modalities have consistently shown brain glucose hypometabolism to be an early irregularity in patients with cognitive impairment and in some cases prelude memory deficits ^1, 11^. Increase in free radical production, a decrease in ATP/ADP ratio and increased rate of oxidant damage to lipids, proteins and mitochondrial DNA characterize the mitochondrial dysfunction in AD ^4, 85^. Whether mitochondrial dysfunction is induced by Aβ or an independent upstream process is currently unresolved. Reduction in expression of dynamin 1 like (*DNM1L*, PEP = 1.4e-03) is consistent with the observation that mitochondrial dynamics shift in favor of fission and is dependent on Aβ overexpression as seen in AD ^4^. Increased fission could also be explained by reduced expression of hypoxia inducible domain family member 1A (*HIGD1A*, PEP = 1.4e-02) shown to regulate mitochondrial fusion by inhibiting the cleavage of *OPA1* ^3^. Interestingly, it has also been attributed to regulate the mitochondrial γ-secretase, a multi-subunit protease complex known to cleave numerous transmembrane proteins including notch and *APP* and whose activity is dysregulated in AD ^36^. Presence of amyloid precursor protein (*APP*) in the mitochondria is associated with reduced cytochrome c oxidase (*COX10*, PEP = 3.8e-05) which is essential for COX assembly and function of complex IV of the electron transport chain ^25^. Although the direct inhibition of COX on Aβ deposition has shown conflicting results in animal and cell culture models ^31, 79^, reduced expression of COX may lead to reduced regulatory heme a key metabolite shown to bind with Aβ ^4^. Remarkably, *HIGD1A* also has a role of positively regulating COX complex presenting with a therapeutic treatment strategy applicable across various disorders with dysfunctional OXPHOS activity including AD ^37, 58^. The expression of malate dehydrogenase 1 (*MDH1*, PEP = 7.0e-03), a key enzyme in the TCA cycle that reversibly catalyzes the oxidation of malate to oxaloacetate was reduced contrary to previous reports of increased protein expression ^4, 85^. This could reflect variance in methodological approaches, patient demography or preferential transcription for this gene. Decreased *MDH1* expression is proposed to enhance oxidative stress and mitochondrial damage due to its interaction with *AKT1* activating a cascade of pathway alterations implicated in AD, PD and ALS ^70^.

Apart from bioenergetics, alteration in metabolic pathways are observed and associated with risk of developing AD. Increased expression of solute carrier family of riboflavin transporters (*SLC52A3*, PEP = 3.6e-02) is intriguing considering riboflavin is a precursor to the complex II substrate flavin adenine dinucleotide. Recent findings of mutations and deficiency of the brain specific transporter SLC52A3 are linked to motor neuron diseases (Brown–Vialetto–Van Laere syndrome, Fazio-Londe Disease) ^12, 60^, while its overexpression was shown to enhance the proliferatory capability of human glioma mediating its effect through suppression of proapoptotic proteins and upregulation of matrix metalloproteinases (MMP) specifically MMP-2 and MMP-9, which have shown to correlate with both CSF Aβ and tau levels from AD patients ^30, 91^. Understanding the signaling pathways of *SLC52A3* warrant further research for validity as clinical diagnostic markers and for dietary considerations ^45^. Adiponectin receptor 1 (*ADIPOR1*, PEP = 8.4e-04) that transduce signals from adiponectin, an adipocyte derived hormone involved in control of fat metabolism and insulin sensitivity was upregulated. Due to its relatively recent discovery, data on *ADIPOR1* gene expression in AD are lacking though increased plasma and CSF adiponectin has been detected in MCI and AD patients ^84^.

### 2.3 Novel regulatory mechanisms of AD pathogenesis include TGIF1 and EGR3

We prioritized transcriptional regulators that were significantly enriched in the AD clustered subnetwork and were differentially expressed along with their gene targets in AD when compared to NDC and identified 4 master transcription factors (TF) i.e. *SP1, EGR3, TGIF1* and *BPTF* across all the clusters (Figure 4, Supplemental Table S8, Figure S6). Specificity protein 1 (*SP1*) is dysregulated in AD and can regulate key genes associated with AD pathology i.e., *APP*, tau and *APOE* ^20^. Gene members of the early growth response family (*EGR3, EGR1)* are zinc finger transcription factors essential for synaptic plasticity and memory and are downregulated. Dysregulation of *EGR1,* a critical microglial homeostatic gene required for maintenance of LTP was associated with pathogenesis of *APP* expressing mice while *EGR3* critical for short term memory has previously not been associated with AD ^50, 65, 69^. *BPTF/ FAC1* is a chromatin remodeler first identified in AD brain whose overexpression is associated with apoptotic cell death ^81^. Transforming growth factor-βs (*TGFβ)* along with bone morphogenic proteins (*BMP)* bind via receptors (*TGFBR1, TGFBR2, TGFBR3, BMPR2)* and activate Smad to regulate gene expression ^64^. Transcriptional co-repressor *TGIF* recruits histone deacetylase (*HDAC*) to modulate Smad complexes to control the *TGF-β* signaling based activation that have pleiotropic functions and are disrupted in AD ^89^.

**Figure 4.**
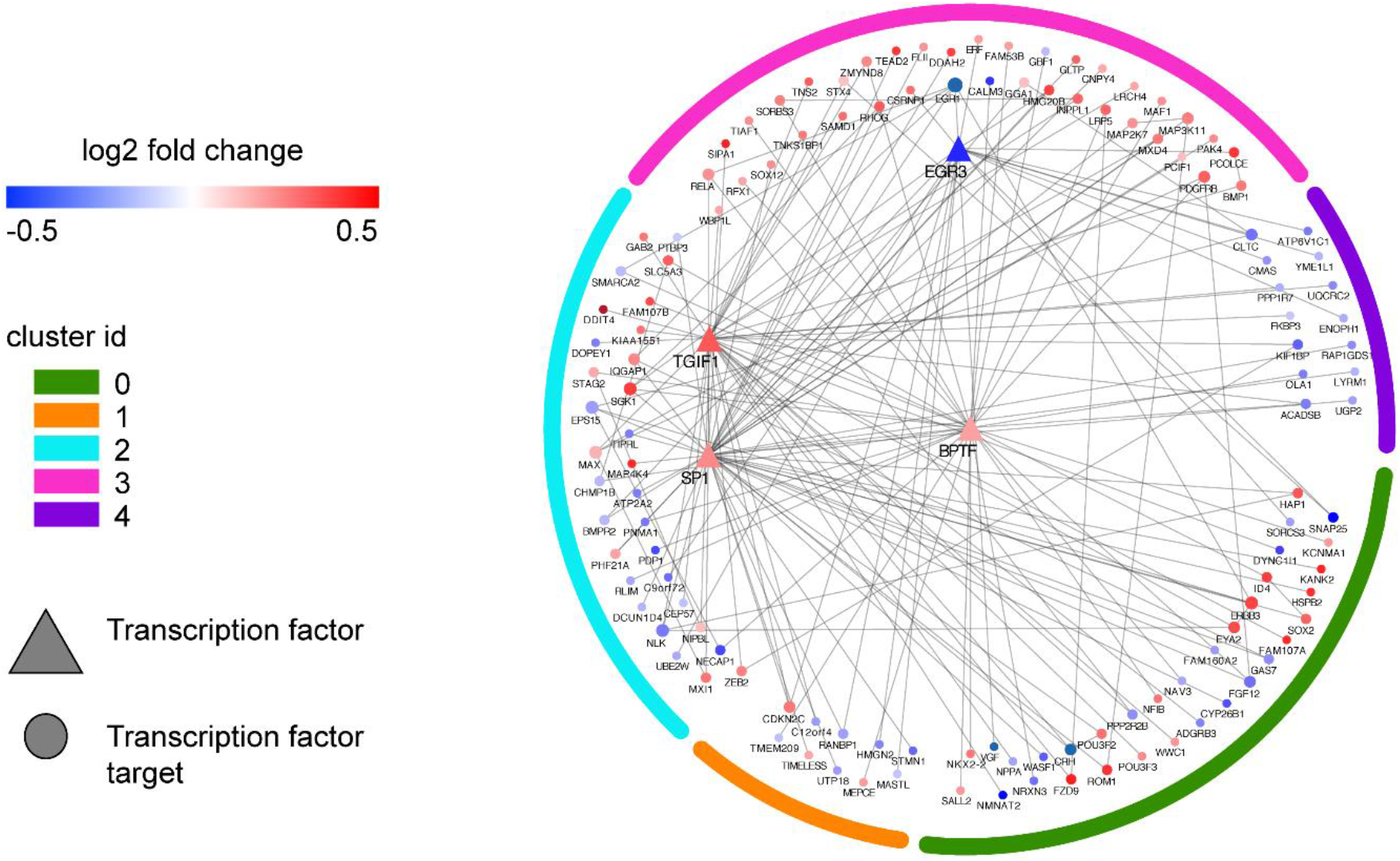

*TGFβ* signaling across multiple cell types can activate several Smad-dependent and independent pathways including MAPKs, NF-κB, PI3K-AKT with contradictory results in a context dependent manner. Consistent with observations in AD brains, overexpression of *TGFβ* and its receptors associated with increased A*β* accumulation along with downregulation of *BMPR2* point towards dysregulation in signaling mediated by *TGFβ* ^89^. It is of interest to note that downregulation of *BMPR2* observed here could be primarily mediated by downregulation of *EGR1,* an essential microglial transcription factor ^65^. Deficiency in *BMPR2* is linked to increased sensitivity to DNA damage along with alterations in DNA repair efficiency and upregulated *SP1* could reflect DNA damage control ^54^. This could provide an important link between neurodegenerative microglial phenotype and DNA damage observed in AD.

Downregulated nemo like kinase (*NLK)* along with upregulation of *NOTCH4* which is shown to strongly couple to AD risk genes and its downstream co-activator complex *RBPJ,* point towards ectopic cell cycle reactivation ^5, 13^. Surprisingly, these observations also coincide with dysregulation of genes involved in reprogramming of cell function i.e. *KLF9* is downregulated while *SOX2, SALL2* and *JUND* are upregulated consistent with reduced efficacy of glucocorticoids (GC) in the aging brain ^47^. Downregulated *KLF9* along with upregulated *JUND* and *SALL2* promote cell proliferation and activate PI3K/AKT pathways involved in growth factors while increased expression of *SOX2, SOX12* can act synergistically with octamer-binding proteins (*POU3F2, POU3F3)* to implement a feed-forward circuit to increase expression of genes (*YES1, ID4)* associated with cell reprogramming. Increased expression of *SGK1,* regulated by *BPTF* could stem from *TGFβ* activated PI3K signaling leading to enhanced activity of NF-κB providing a positive feedback by cross talk between the two pathways ^47, 64^.

#### 2.3.1 TGIF1 is neuroprotective in AD

Here we propose increased *TGIF1* plays a neuroprotective role by directly repressing or by activating co-repressors to regulate gene expression aimed at arresting cell cycle, enhancing DNA repair and restoring homeostasis (Figure 5). Reduced expression of glycogen synthase kinase 3 B (*GSK3B)* results in prolonged activation of Smad by inhibition of proteasome-mediated degradation. Increased expression of DNA-binding transcriptional repressor *ZEB2* targets activated Smads, which are hub proteins for pathway regulation and control expression for diverse set of genes ^64^. *WWC1* and its protein product KIBRA which are associated with AD, are essential for hippo signaling and synaptic plasticity, with increased expression resulting in inhibition of proliferation ^48^. Upregulation of negative regulator of *MYC* max interactor 1 (*MXI1*) and transcription regulator myc associated factor x (*MAX*) point towards inhibition of *MYC* which drives cell cycle re-entry and is increased in AD brains ^10^. Tensin 2 (TNS2) is a focal adhesion protein that regulates signaling pathways for cell shape and motility by bridging cytoskeleton and intracellular projections of transmembrane receptors. By acting as a protein tyrosine phosphatase, *TNS2* dephosphorylates *IRS1* reducing its activation for binding to signaling molecules and reduces Akt signaling via a negative feedback loop ^39^. Centrosomal protein 57 (*CEP57*) mediates the mitogenic activity of fibroblast growth factor 2 (*FGF2*) which shows enhanced binding to senile plaques and neurofibrillary tangles and stimulates downstream signaling pathways (MAPK/ERK, PLC, Akt) ^22, 47, 75^. Together with the upregulation of fibroblast growth factor receptor 2 (*FGFR2*), *FGF2* possibly potentiates glial mediated neuroinflammation in AD and *TGIF1* acts on *CEP57* to lead to a net suppression effect on signaling mediated by *FGF2* ^52^. Eyes absent proteins (*EYA2*) are transcriptional cofactors with an intrinsic phosphatase activity which phosphorylates histone H2AX to mediate DNA repair from double strand breaks ^51^. Interestingly, upregulation of *DDIT4,* an inhibitor of mTOR activity which was found to be hyperactivated in AD and mild cognitive impairment but not in pathological aged brains is consistent with role of *TGIF1* as neuroprotective ^86^.

**Figure 5.**
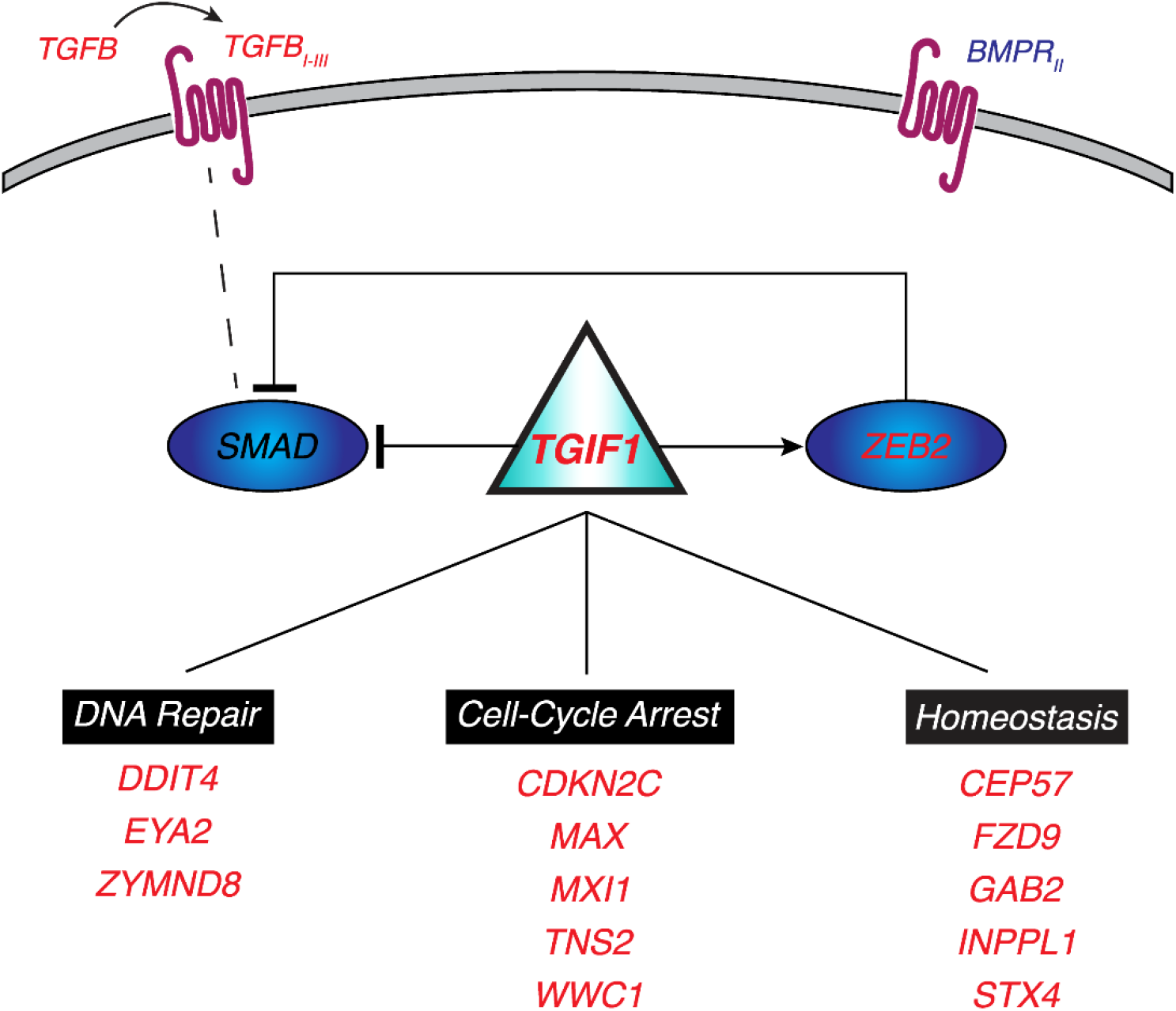

Upregulation of grb-associated binding protein 2 (*GAB2)*, a consistently replicated GWAS identified AD gene is considered to be neuroprotective and correlates with decreased neurofibrillary tangles and senile plaques ^101^. Some of the *TGIF1* regulated genes have implication for synaptic transmission. Increased expression of *INPPL1* together with decreased *SYNJ1* and increased *AKT1* could be interpreted as compensatory mechanism to restore the imbalance in the phosphoinositide pool which is crucial for synaptic function and to regulate PI3K/Akt signaling ^8^. Similarly, increased expression of syntaxin 4 (*STX4*) critical for vesicle docking, Wnt frizzled receptor (*FZD9*) along with co-receptor (*LRP5*) which regulate synaptic vesicle cycle may be rescue aimed to compensate for effects mediated by *EGR3* (see below) ^42^.

*TGIF1* also cooperates with the other transcription regulators to mediate the neuroprotective effect (Supplemental Figure S7). Upregulation of *CDKN2C* regulated by *TGIF1* and *EGR3* is perhaps a compensatory effect to inhibit the cell cycle progression in absence of direct regulation by *TGFβ* ^64^. *BPTF* and *TGIF1* regulated zinc-finger mynd-type containing 8 (*ZMYND8*) is a chromatin reader that recruits the repressive nucleosome remodeling and histone deacetylase (NuRD) complex to transcriptionally silence gene expression as part of the DNA damage response. In conjunction with increased expression of *BRD3*, paralog of *BRD2*, Z*MYND8* in AD may work to shield H4Ac during double strand repairs as part of the chromatin response to DNA damage ^35^.

#### 2.3.2 EGR3 affects synaptic vesicle processing in AD

Consistent with the association of synaptic loss to AD, we observe a concerted downregulation of many genes that code for proteins involved in synaptic vesicle cycle of which key genes are regulated by *EGR3* (Figure 6). Vacuolar ATPase (V-ATPase) is an essential proton pump highly expressed on the membrane of the presynaptic vesicle, facilitates neurotransmitter concentration in synaptic vesicles and are differentially expressed in brains of different *APOE* isoforms ^96^. C1 subunit of V-ATPase (*ATP6V1C1*) is crucial to connect the ATP catalytic domain (V_1_) to the proton-translocation domain (V_0_) and its downregulation has implications on vesicle acidification as well as on membrane fusion. Reduction in expression of neuronal SNARE synaptosome associated protein 25 (*SNAP25*) along with synaptotagmins (*SYT5, SYT13)* and syntaxin-binding proteins (*STXBP1, STXBP5L*) indicate reduced vesicle docking and SNARE-mediated fusion ^7^. Furthermore, decrease in vesicle-fusion ATPase (*NSF)* necessary for disassociating SNARE complex from the plasma membrane implicates decreased availability of uncomplexed-SNARE ^43^. Reduced clathrin (*CLTC)* along with adaptor protein complex (*AP2A*), synaptophysin (*SYP*) and rab (*RAB3A*) imply reduced vesicle recycling ^7^. Targeting the regulation of the dysfunction of clathrin-mediated endocytosis for synaptic receptor holds promise for early intervention strategy ^78^. Neurosecretory polypeptide precursor *VGF* is enhanced in neuronal activity associated with LTP and can be postulated to have a role in sorting of other regulated secretory proteins including glutamate in immature vesicles ^38^. Sortilin-related receptor (*SORCS3*), a neuronal receptor for proteolytic processing of *APP* is implicated in AD and is shown to modulate LTD ^14^. *EGR3* mediated reductions in *VGF* and *SORCS3* aid in the observed phenotype of Aβ induced impairment in LTP and neuronal networks ^78^.

**Figure 6.**
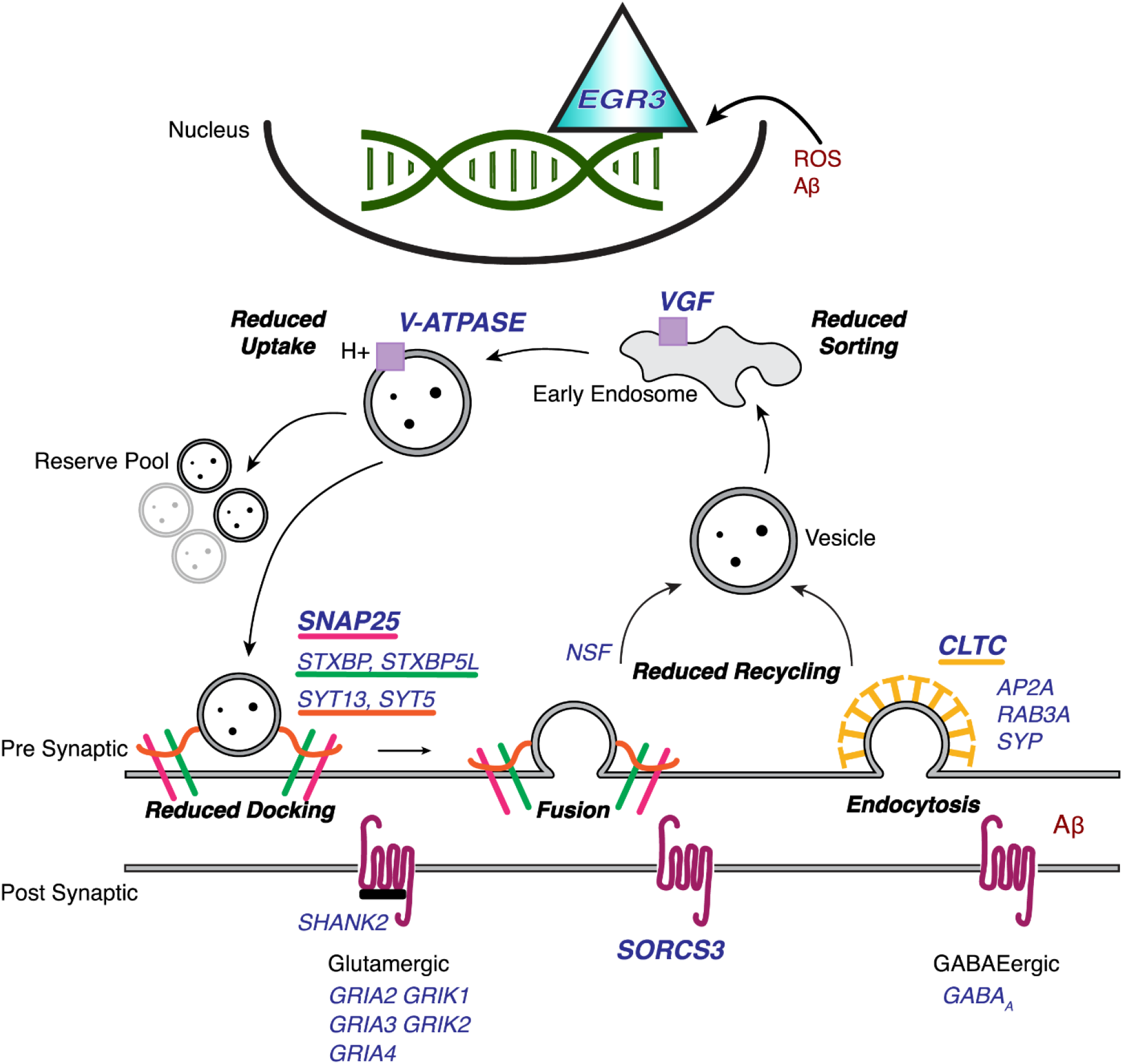

### 2.4 Protein expression of TGIF1 and EGR3 is in concordance with transcript level expression

To substantiate the differences in transcription between AD and non-demented aged brains, we assessed the protein expression of *TGIF1* and *EGR1,* the direct target for *EGR3* in pre-frontal cortex of AD and age-matched control brains from an independent cohort. In line with changes in the transcriptome data, we observe an overall 60% decrease in *EGR1* and an 81% increase in *TGIF1* levels in AD brains relative to non-demented control and these trends reach statistical significance. (Figure 7). We found the *EGR1* distribution across all cell types in the control brains and the protein levels were particularly diminished in pyramidal neurons in AD brains. On the other hand, upregulation of the nuclear localization of *TGIF1* was a frequent feature of glial cells in AD brains. Taken together, these results demonstrate the downregulation of *EGR3* and is consistent with the role of *TGIF1* as a transcription factor.

**Figure 7.**
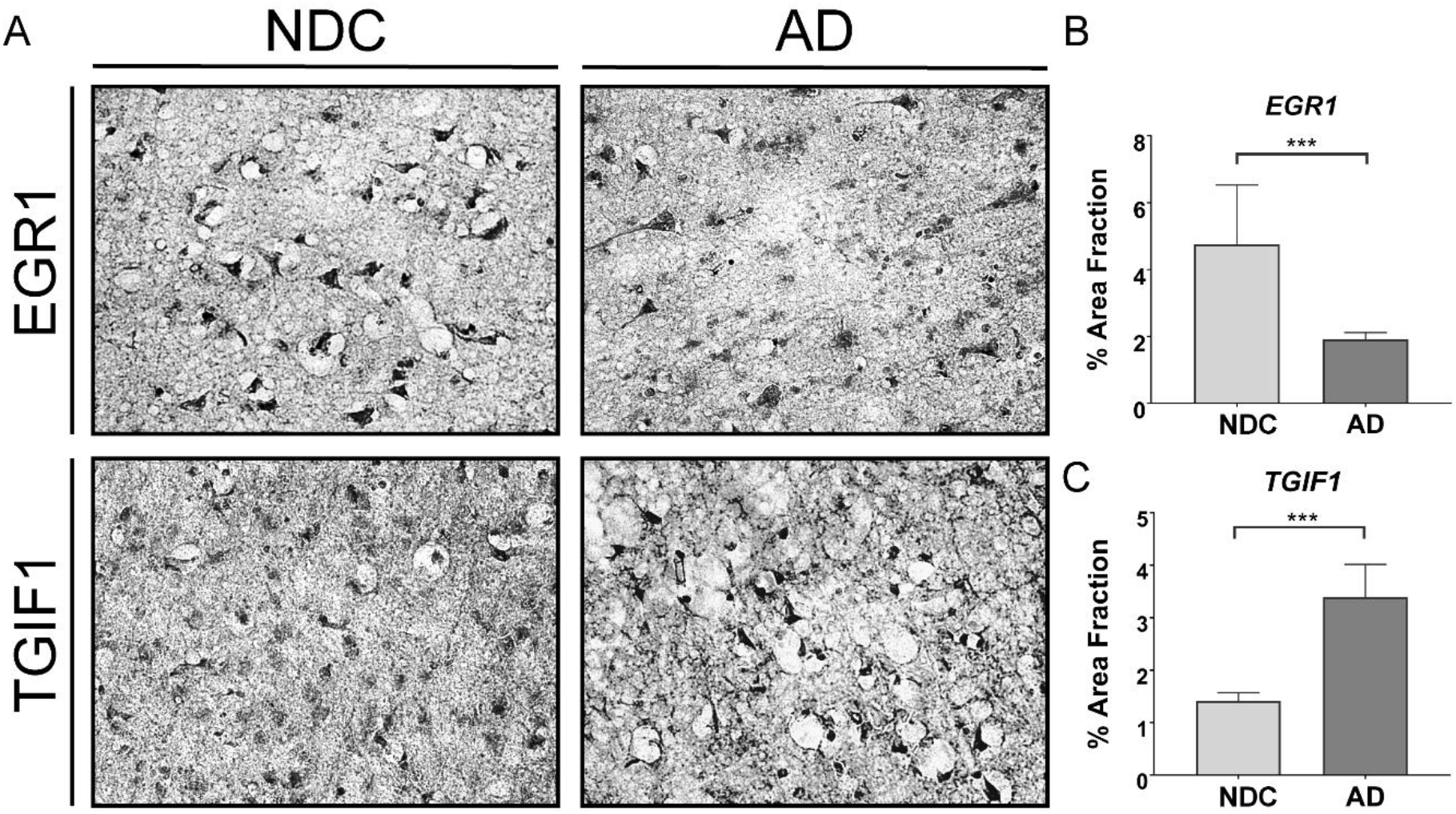

## 3. Conclusions

By integrating brain tissue specific protein interactions with gene expression profiles derived from large sampling of postmortem transcriptome data, we identified changes in molecular network associated with clinical and pathological Alzheimer’s disease (AD) when compared to age matched non-demented controls. Although the mechanisms for neuronal dysfunction in AD are not completely understood yet, the distinct biological clusters identified using Louvain community detection algorithm point towards associations critical for AD ^9, 23, 46, 68, 85, 99^. Interestingly, the average global expression for clusters corresponding to synaptic transmission, metabolism, cell cycle and survival as well as immune response were downregulated while the upregulated cluster3 had a large set of uncharacterized pathways and processes. By refining the subject selection, we identified new genes with target potential, confirmed some previous correlations and find lack of associations for other previously reported GWAS computed AD susceptible genes. Based on gene interactions, we propose neuroprotective role for transcriptional repressor *TGIF1* which activates other co-repressors to module the *TGFβ* signaling and acts to arrest cell cycle re-entry. We find downregulation of *EGR3* to be associated to dysregulated synaptic vesicle cycle and postulate to mediate synaptic deficits as seen in AD.

## 4. Methods

### 4.1 Study Subjects

The Religious Orders Study and Rush Memory and Aging Project (ROSMAP) are an ongoing longitudinal clinical-pathologic cohort studies of aging and dementia with enrollment of aged individuals without known dementia at baseline ^6^. All participants are organ donors. Participants undergo detailed annual clinical evaluation. At death, a postmortem brain evaluation is performed, including silver stain to assay AD pathology (neuritic and diffuse plaques, and neurofibrillary tangles), and Aβ load by image analysis and the density of PHF tau-positive neurofibrillary tangles ^6^. The Institutional Review Board of the Rush University Medical Center approved the both studies. An informed consent and an anatomical gift act are obtained from each participant, as is a repository consent that allows for sharing of data and biospecimens.

Subjects were classified as either non demented controls (NDC) or AD based on a weighted combination of complementary information including cognitive score (MMSE) within two years of death along with three separate pathological scores i.e., total amyloid load, neurofibrillary tangles (Braak stage), and presence of neuritic plaques (CERAD score) ^41^. Non-demented controls (NDC) included cognitively intact subjects (no diagnosis of mild cognitive impairment within two years of death) containing diffuse amyloid deposits in absence of neuritic plaques, and neurofibrillary tangles confined to the entorhinal region of the brain. For subjects with cognitive impairment, AD diagnosis was assigned based on the high or intermediate likelihood of AD neuropathology as presented in the revised NIA-AA guidelines for neuropathologic assessment of disorders of the brain common in the elderly ^41^. A total of 414 subjects who passed these criteria were included in the current study (Table 1). Assessment of the pathological metrics as well as the characteristics of the cohort has been previously described in detail ^6, 95^.

**Table 1:** Sample demographics of the subjects included in the study.

| **Demographic Varle** | **Non Demented Controls (NDC) (N = 151)** | **Alzheimer's Disease (AD) (N = 263)** |
| --- | --- | --- |
| Female Sex, No. (%) | 92 (60.9) | 185 (70.3) |
| Education, years | 16.5 (3.6) | 16.2 (3.5) |
| MMSE | 28.6 (1.0) | 14.5 (7.5) |
| *APOE* $\epsilon^4$, No. (%) | 20 (13.2) | 90 (34.2) |
| Age at Death, years | 85.2 (6.6) | 90.9 (5.9) |
| PMI, hours | 7.4 (4.4) | 7.6 (5.2) |
| RIN | 7.2 (1.0) | 6.9 (0.9) |

Abbreviations: AD, Alzheimer's Disease; APOE, apolipoprotein E; MMSE; Mini-Mental State Examination; PMI, Postmortem Interval; RIN, RNA Integrity Number
Data presented as Mean (S.D) unless specified

### 4.2 Gene Expression Analysis

RNA-seq data (101 bp paired end reads with coverage of 50 million) was obtained from RNA extracted from the gray matter of the dorsolateral prefrontal cortex. Expression in the form of fragments per kilobase of exon per million reads mapped (FPKM) was estimated by RSEM. Details of the RNA-seq data pipeline can be found in previous literature ^68, 95^.

Expression data were downloaded from ROSMAP as FPKM values and normalized using a multi-loess method and batch effects removed using COMBAT algorithm ^44^. For the log-transformed expression data, linear regression models were fit to account for effects of diagnostic conditions as well as confounding covariates of age at death, sex, postmortem interval (PMI) and APOE status for the risk allele. Test for statistical significance was achieved by implementation of a Bayesian strategy of Lönnstedt and Speed as implemented in *R* package *limma* ^72^. Genes are sorted by their posterior error probability (PEP) and considered significant at PEP < 0.05.

### 4.3 Background Interactome

The background interactome on which the network analysis was conducted was built from the brain-specific network in the GIANT database ^34^. This network is composed of edges which support a strong tissue-specific functional interaction between source and target genes. This network, thresholded at an edge confidence of 0.2, contains 14,545 genes, and 1,370,174 edges, and represents genes and interactions which are likely to be present in normal brain function. The identified differentially expressed genes were mapped onto this network to create an AD subnetwork. Network visualization was accomplished using visJS2Jupyter ^73^.

### 4.4 Clustering analysis

Clusters were identified in the AD subnetwork using a network-based modularity maximization algorithm ^88^. This algorithm, commonly referred to as the Louvain clustering algorithm, identifies groups of nodes which have many connections within groups, and few connections between groups, and is efficient at extracting community structure from large networks. The algorithm iteratively maximizes the modularity, Q, defined as follows:

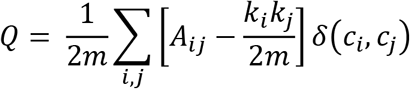

Where *A*_*ij*_ is the binary adjacency matrix element representing the presence or absence of the connection between node *i* and node *j, k_i_* represents the degree of node *i*, where degree is defined as the number of nodes directly connected to node *i*, *c*_*i*_ is the community to which node *i* belongs, and the function *δ*(*x*, *y*) is 1 if *x* = *y*, and otherwise it is 0. *m* is the total number of edges in the network, 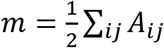 (divided by two to avoid double counting). See Blondel et al ^88^ for more details on the algorithm.

The Louvain algorithm does not require selection or fine-tuning of parameters, as do some other clustering algorithms such as, K-means clustering and hierarchical clustering (such as are used in the WGCNA R package). Rather, clusters are determined by finding groupings of genes which have many within group connections, and few between group connections. This algorithm has proven useful in detecting modules, or clusters, in protein-protein interaction networks, such as the one used here ^29, 87^.

### 4.5 Functional Annotation and Pathway Analysis

To identify the underlying biological functions enriched in AD, Gene Set Enrichment Analysis (GSEA) was implemented which identifies the enrichment of functionally defined gene sets using a modified Kolmogorov-Smirnov statistic ^82^. The Molecular Signature Database (MSigDb v6.0) ^82^ consists of well over 4000 gene sets from functionally well-established or published pathways from databases including Gene Ontology (GO), the Kyoto Encyclopedia of Gene and Genomes (KEGG), BioCarta, and Reactome. Statistical significance after adjusting for multiple testing is defined at FDR < 0.05. Gene set-based permutation test of 1000 permutations was applied. Detailed description of the GSEA algorithm, testing metrics and the implementation have been described previously ^82^. Hypergeometric test was utilized to test of statistical significance of the enriched biological process and pathways identified for the differential expressed genes in the clusters ^90, 97^. Overrepresentation enrichment analysis was conducted using the full set of genes from the AD subnetwork as the reference gene set, corrected for multiple testing using the Benjamini-Hochberg procedure and FDR < 0.05 was considered significant.

### 4.6 Transcriptional Regulation Analysis

To identify transcription factors (TF) likely related to AD, we analyzed for enrichment of transcription factor targets in the AD clustered subnetwork ^90^. This resulted in 16 significantly enriched transcription factors (Supplementary Table S8). We further filtered this list by differential expression in AD vs NDC, resulting in 4 candidate TFs. These four TFs are all significantly differentially expressed in AD as compared to NDC and have more targets than would be expected by chance in the AD subnetwork (Supplemental Figure S7). The TF subnetwork is visualized to highlight the connections between TF and targets along with connections between targets using visJS2Jupyter ^73^.

### 4.7 Validation of *TGIF1* and *EGR3*

We assessed the protein expression and distribution of *TGIF1* and *EGR3* target *EGR1* in prefrontal cortex brain tissue of clinically diagnosed pathologically confirmed AD and from age matched controls using standard immunohistochemistry techniques. The subjects were selected based on diagnostic criteria as described above and brain tissue samples were obtained from University of California, San Diego Alzheimer’s Disease Research Center (ADRC) brain bank. Tissue samples were obtained from two non-demented aged controls (81.5 ± 2.1 years, 12 ± 3 hours, 50% F) and three AD subjects (83 ± 4.3 years, 6.6 ± 1.1 hours, 67% F). Paraffin-embedded tissue blocks were serially sectioned and incubated with either TGIF antibody (H-1) (Santa Cruz Biotechnology, Dallas, TX) or EGR1 antibody (Cell Signaling Technology, Danvers, MA). Staining was performed with chromogen 3,3^’^ -diaminobenzidine (DAB) to identify the immunoreactive structures and counterstained with hematoxylin. All images were acquired using an upright microscope (Leica DM5500B) at a resolution of 1392 x 1040 pixels and consistent aperture and gain settings. Custom designed macro was used to convert the optical images to grayscale, threshold and measure the TGIF or EGR1 positive area fraction relative to the optical field. Due to the non-normal distribution of the data, test of significance was evaluated using Wilcoxon-Mann-Whitney test and differences were considered significant when p < 0.05.

## Acknowledgements

We gratefully acknowledge the study participants. This study was funding by the VA San Diego, Grant BX003040. The contents do not represent the views of the U.S Department of veterans Affairs or the United States Government. The project described was partially supported by the National Institutes of Health, Grant UL1TR001442 of CTSA, and P30AG10161, RF1AG15819, R01AG17917, U01AG46’52. The content is solely the responsibility of the authors and does not necessarily represent the official views of the NIH.

